# *RET* enhancer haplotype-dependent remodeling of the human fetal gut development program

**DOI:** 10.1101/2022.07.26.501565

**Authors:** Sumantra Chatterjee, Lauren E. Fries, Or Yaacov, Nan Hu, Hanna E. Berk-Rauch, Aravinda Chakravarti

## Abstract

Hirschsprung disease (HSCR) is associated with deficiency of the receptor tyrosine kinase RET, resulting in loss of cells of the enteric nervous system (ENS) during fetal gut development. The major contribution to HSCR risk is from common sequence variants in *RET* enhancers with additional risk from rare coding variants in many genes. Here, we demonstrate that these *RET* enhancer variants specifically alter the human fetal gut development program through significant decreases in gene expression of *RET*, members of the *RET-EDNRB* gene regulatory network (GRN), other HSCR genes, and an altered transcriptome with 2,382 differentially expressed genes with diverse neuronal and mesenchymal functions. A parsimonious hypothesis for these results is that beyond RET’s direct effect on its GRN, it also has a major role in enteric neural crest derived cell (ENCDC) precursor proliferation, its deficiency reducing ENCDCs with relative expansion of non-ENCDC cells. Thus, genes reducing RET proliferative activity can potentially cause HSCR. One such class is the 23 *RET*-dependent transcription factors enriched in early gut development. We show that their knockdown in human neuroblastoma SK-N-SH cells reduces *RET* and/or *EDNRB* gene expression, expanding the *RET-EDNRB* GRN. The human embryos we studied had major remodeling of the gut transcriptional but were unlikely to have had HSCR: thus, genetic changes in addition to those in *RET* are required for a significant enough reduction in ENCDCs to cause aganglionosis.

## Introduction

Studying the spatial and temporal genetic program of human tissue development, and its alterations in disease, are challenging primarily owing to the difficulty of gaining access to the relevant tissues during development. Much progress has been made by extrapolation from cognate studies of wildtype and gene mutations in model organisms (Chatterjee, Sivakamasundari et al. 2014, Chatterjee, Nandakumar et al. 2019) but this approach cannot replace direct studies of human organogenesis. Here, we demonstrate how we can gain mechanistic insight into genetic programs of normal versus compromised human development using functional genotypes common in humans.

An exemplar of this approach is Hirschsprung disease (HSCR, congenital aganglionosis), characterized by the absence of enteric ganglia along variable lengths of the distal colon with an absence of gut motility (Chakravarti and Lyonnet 2001). This developmental defect of the enteric nervous system (ENS) arises from the failure of ENCDC precursors to differentiate, proliferate and migrate in the gut (Obermayr, Hotta et al. 2013). HSCR is associated with rare coding pathogenic alleles (PAs) in 24 genes (Tilghman, Ling et al. 2019), as well as common non-coding variants at *RET* (Emison, McCallion et al. 2005, Emison, Garcia-Barcelo et al. 2010, Chatterjee, Kapoor et al. 2016), *NRG1* (Garcia-Barcelo, Tang et al. 2009) and *SEMA3C/D* (Jiang, Arnold et al. 2015). Rarer chromosomal aberrations and large copy number variants (CNVs) also make significant contributions to risk (Tilghman, Ling et al. 2019). Of all genes, the receptor tyrosine kinase gene *RET* is key to the ENS developmental program because ∼50% of HSCR patients carry *RET* PAs, the vast *majority* of which are rare partial or total loss-of-function coding mutations (Tilghman, Ling et al. 2019). However, the greatest risk contribution to HSCR arises from multiple common variants within transcriptional enhancers of *RET* (Chatterjee, Kapoor et al. 2016, Chatterjee, Karasaki et al. 2021). The high frequency of these hypomorphic *RET* enhancer variant genotypes thus enable direct comparisons of their genetic programs in the developing gut in randomly collected human embryonic specimens.

There are many aspects of human gut neurogenesis, a process initiated at Carnegie stage (CS) 14 (week 4 of gestation) and completed by CS22 (week 7) (Goldstein, Hofstra et al. 2013), that are unknown. During this interval, a mass of undifferentiated mesoderm organizes under inductive signals to eventually form the layers of the gut and to establish innervation, circulation and immune functions. Disruptions in these processes can lead to a dysfunctional ENS and enteric neuropathies manifesting with abnormal gut motor function. The classic enteric neuropathy is HSCR which is the most common cause of functional obstruction of the neonatal gut. Despite clues to its genetic origins, and the major effect of *RET*, it is still unknown how its pathogenic variants lead to aganglionosis. Further, what is common to the diverse genetic defects in HSCR that they all lead to aganglionosis?

We attempt to answer these questions using three non-coding risk variants at *RET*, rs2506030, rs7069590, and rs2435357, that are highly polymorphic and reside within three enhancers (RET-7, RET-5.5, RET+3) bound by the transcription factors (TFs) RARB, GATA2 and SOX10 (Kapoor, Jiang et al. 2015, Chatterjee, Kapoor et al. 2016). Recently, we have identified additional *RET* enhancers also with common HSCR-associated variants, at least two of which bind PAX3 (Chatterjee, Karasaki et al. 2021). Nevertheless, a significant portion of the noncoding risk at *RET* is from a haplotype (S, susceptible) comprising the three risk alleles rs2506030, rs7069590, and rs2435357; the S haplotype has significantly lower *RET* gene expression in human fetal guts as compared to the complementary (R, resistant) haplotype (Chatterjee, Kapoor et al. 2016, Chatterjee and Chakravarti 2019). Further, *in vitro* deletion of these three enhancers in the human neuroblastoma SK-N-SH cell line leads to loss of *RET* expression, providing evidence of their direct role in *RET* regulation (Chatterjee, Karasaki et al. 2020). We use these common enhancer genotypes, RR, RS and SS, to stratify human fetal tissues and study their effects on gut neurogenesis. Although such expression quantitative trait loci are routinely studied in accessible adult tissues (Cheung and Spielman 2002, Spielman, Bastone et al. 2007, Consortium 2013, Kim-Hellmuth, Bechheim et al. 2017, Kim-Hellmuth, Aguet et al. 2020), and gene expression atlases of human embryonic tissue have been successfully produced (Gerrard, Berry et al. 2016, Cardoso-Moreira, Halbert et al. 2019, Groff, Resetkova et al. 2019), there is very limited work on the consequences of disease-associated variation on fetal development (Jaffe, Straub et al. 2018, O’Brien, Hannon et al. 2018).

In this study, we build global gene expression maps of 23 human fetal gut samples at CS14 and CS22, for the RR, RS and SS *RET* enhancer genotypes, to demonstrate the profound ways in which *RET* modulates the ENS genetic program. We demonstrate (1) the gradual loss of *RET* expression with increasing S haplotype dosage; (2) significant changes in expression of *RET-EDNRB* gene regulatory network (GRN) genes; (3) significant changes in expression of the majority of known HSCR genes; (4) expansion of the *RET-EDNRB* GRN to many TFs that not only regulate *RET* and *EDNRB* but are also under *RET* feedback control; (5) increasing transcriptomic dysregulation of neurogenesis, cell cycle regulation and signal transduction pathways; and, (6) non-cell autonomous effects of RET on extra-cellular matrix (ECM) formation. These studies point to a broader role of *RET* during gut morphogenesis than previously envisioned and the many molecular processes compromised in *RET* deficiency, suggesting causes of HSCR clinical phenotypes beyond aganglionosis as well as novel gene targets for mutational analysis in HSCR. These analyses are easily envisioned for genes and tissues in other developmental disorders.

## Results

### RET-dependent gene expression changes in the developing human gut

We obtained 23 (3 at CS14 and 20 at CS22) human fetal gut (stomach, foregut and hindgut) tissues from the Human Developmental Biology Resource (HDBR) (Gerrelli, Lisgo et al. 2015) and genotyped them for three HSCR-associated *RET* enhancer polymorphisms (rs2506030 (A/**G**), rs7069590 (C/**T**), rs2435357 (C/**T**); risk alleles in bold) to classify them by their resistant (R: **GT**C, A**T**C, **G**CC, ACC) or susceptible (S: A**TT, GTT**) haplotypes (Chatterjee, Kapoor et al. 2016). Our sample comprised 2 RR and 1 RS genotype at CS14, and 9, 8, and 3 embryos with RR, RS, and SS genotypes at CS22 (**Table 1**). Tissue-level expression profiling by RNA-seq on these 23 samples revealed no global gene expression differences among them, with mean normalized read counts (log2 scale) of 7.39± 3.36, 7.38± 3.33 and 7.41± 3.31 for RR, RS and SS, respectively (P= 0.37) (**Figure 1A**). At CS14, the corresponding figures for RR and RS were 5.52± 2.71 and 5.51± 2.73, highlighting an overall increase in gut gene expression over developmental time irrespective of genotype.

**Table 1:** Genotypes of 3 Hirschsprung disease associated polymorphisms (rs2506030, rs7069590 and rs2435357) in 20 fetal gut samples at Carnegie stage (CS) 22 and 3 samples at CS14. Risk alleles are marked in bold for each polymorphism. **R** is resistant and **S** is susceptible haplotypes.

| Sample ID | rs2506030 | rs7069590 | rs2435357 | Haplotype |
| --- | --- | --- | --- | --- |
| **CS22** |  |  |  |  |
| HFG7 | **AG** | CT | CC | R/R |
| HFG10 | **AG** | **TT** | **TT** | S/S |
| HFG14 | AA | **TT** | CC | R/R |
| HFG17 | **AG** | **TT** | CT | R/S |
| HFG20 | AA | **TT** | CC | R/R |
| HFG21 | **AG** | **TT** | CC | R/R |
| HFG22 | **GG** | **TT** | **TT** | S/S |
| HFG23 | AA | **TT** | CT | R/S |
| HFG24 | **AG** | **TT** | CC | R/R |
| HFG25 | **AG** | **TT** | CC | R/R |
| HFG31 | **GG** | **TT** | CT | R/S |
| HFG33 | **AG** | CT | CT | R/S |
| HFG35 | **AG** | **TT** | CT | R/S |
| HFG36 | **AG** | CT | CT | R/S |
| HFG37 | AA | **TT** | CC | R/R |
| HFG39 | **AG** | **TT** | CC | R/R |
| HFG41 | AA | **TT** | CT | R/S |
| HFG42 | **AG** | CT | CT | R/S |
| HFG43 | **GG** | **TT** | **TT** | S/S |
| HFG44 | AA | CT | CC | R/R |
| **CS14** |  |  |  |  |
| HFG47 | **AG** | CT | CC | R/R |
| HFG48 | AA | **TT** | CT | R/S |
| HFG49 | **AG** | **TT** | CC | R/R |

**Figure 1:**
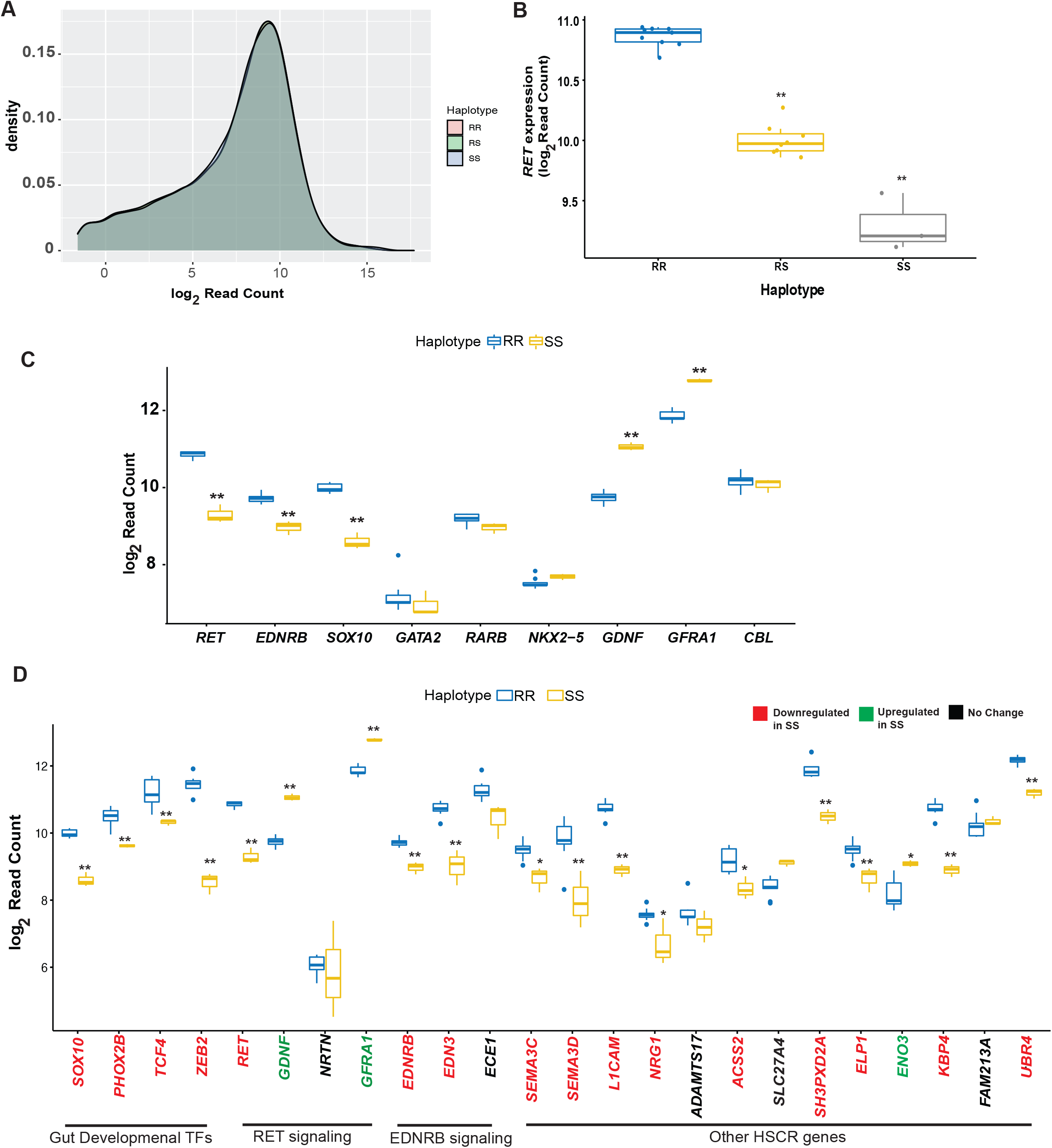
*RET* haplotype-dependent gene expression of Hirschsprung disease genes in the Carnegie Stage 22 (CS22) human fetal gut. (A) Transcriptomes of 20 human fetal guts at CS22 classified by *RET* genotype, based on resistant (R) and susceptible (S) enhancer haplotypes, demonstrate no global differences in their expression profiles (P = 0.37). (B) *RET* gene expression in the developing fetal gut is exponentially reduced with increasing dosage of the S haplotype. (C) Gene expression of 9 genes comprising the *RET-EDNRB* gene regulatory network identifies 5 members with significant RR versus SS differences. (D) Gene expression differences between RR and SS genotypes at 24 HSCR genes: *NRTN, ECE1, ADAMTS17, SLC27A4* and *FAM213A* have no significant change in expression; *GDNF, GFRA1* and *ENO3* are upregulated while the remaining 16 are downregulated in *RET* deficient (SS) embryonic guts. (*, **: Benjamini-Hochberg FDR < 0.01, 0.001).

*RET* gene expression was, however, significantly different between haplotypes (**Figure 1B**). At CS14, RR and RS genotypes had mean log2 read counts of 7.15 and 6.04, respectively, showing a 2.1-fold decrease (P = 0.0012) with one S haplotype; at CS22, there is significant loss of *RET* expression with increasing S haplotype dosage, namely, mean log2 read counts were 10.86, 10.02 and 9.29, which are significantly different between RR and RS (1.8-fold, P = 4.68×10^−10^) as well as between RS and SS (1.6-fold, P = 10^−6^). SS guts have 3-fold lower *RET* expression compared to RR (P=4.65×10^−12^) (**Figure 1B**). This enhancer haplotype is a highly significant eQTL regulating *RET* with the S haplotype being expression deficient relative to the R haplotype in the developing tissue of interest. This is direct evidence that the association between *RET* enhancer genotypes and HSCR arises from loss of *RET* gene expression in the developing gut, analogous to *RET* coding mutations (Kjaer, Hanrahan et al. 2010)

### RET-dependent RET-EDNRB GRN gene expression changes in the developing human gut

We have previously demonstrated that some HSCR genes are not transcriptionally independent of *RET* but united through a GRN controlling *RET* and *EDNRB* gene expression. Through this GRN, *RET* and *EDNRB* also exert feedback regulatory control on other GRN members, including its TFs. Our previous studies of *RET* knockdown in human SK-N-SH neuroblastoma cells (Chatterjee and Chakravarti, 2019), which expresses all known members of the *RET-EDNRB* GRN and shows ligand (GDNF)-dependent RET activation, and in mice with a *RET* LoF mutation demonstrated that many genes within this GRN are transcriptionally affected by complete RET deficiency (Kapoor, Auer et al. 2017, Chatterjee, Nandakumar et al. 2019). These results are recapitulated in the developing human gut in SS genotypes in the human *RET-EDNRB* GRN (**Figure 1C**). At CS22 we observe transcriptional upregulation of the *RET* ligand GDNF and co-receptor GFRA1, and downregulation of its TF SOX10. Further, we observe transcriptional downregulation of *EDNRB* in the SS gut as well.

There is, however, no significant transcriptional change in the three other *RET* TFs, GATA2, RARB and NKX2-*5*, whereas a fifth TF PAX3 is not expressed at this developmental stage. There is also no significant expression change in *CBL*, the ubiquitin ligase targeting phosphorylated RET (**Figure 1C**). These results at CS22 are fully consistent with our studies in the E14.5 *Ret* null mouse, the equivalent stage of mouse gut neurogenesis, where *Gata2, Rarb, Nkx2-5* and *Cbl* are also unaffected (Chatterjee, Nandakumar et al. 2019).

### Effect of RET on HSCR disease genes

Our data can be used to ask whether other HSCR genes, not currently known to be members of the *RET-EDNRB* GRN, are also affected by *RET* deficiency. Consequently, we asked whether all 24 known HSCR disease genes (Alves, Sribudiani et al. 2013, Tilghman, Ling et al. 2019) are *RET*-dependent, implying that they are part of the same GRN. This was tested by association between gene expression levels of these 24 genes, all of which are expressed in the CS22 gut, with S haplotype dosage. Sixteen HSCR genes are significantly (FDR<0.001) downregulated, *GDNF, GFRA1* and *ENO3* are upregulated, while *NRTN, ECE1, ADAMTS17, SLC27A4* and *FAM213A* show no difference (**Figure 1D)**. Therefore, in HSCR, *RET* deficiency can have its effect amplified by altering gene expression of many members of the HSCR gene universe, highlighting the transcriptional connectivity of seemingly functionally unrelated genes leading to the same disease.

### Global effects of RET-dependent gene expression changes in the developing human gut

Beyond the effect on HSCR related genes, we detected 2,382 differentially expressed (FDR<0.01) genes between the transcriptomic profiles of RR and SS: 1,655 genes were downregulated and 727 upregulated in SS (**Figure 2A**). Similar analysis comparing RS with SS identified 2,334 differentially expressed genes (1,631 downregulated and 703 upregulated); in contrast, RR and RS identified 33 differentially expressed genes (28 downregulated and 5 upregulated). Hence, loss of *RET* expression with two S haplotypes is necessary to generate a major effect by altering 8% of the developing gut transcriptome with the majority (69%) of genes showing decreased gene expression. Thus, in normal gut development, *RET* acts to activate gene expression, as also observed in the mouse, where at E14.5, a comparable developmental stage, there are 325 down-regulated and 111 up-regulated genes (Chatterjee, Nandakumar et al. 2019). The large difference between the mouse and human embryonic genes affected is perhaps not unexpected given the single gene difference in the mice compared to the millions of common variants differing between the human samples. Nevertheless, this is a surprisingly large effect because *RET* is not a direct activator or repressor of transcription. To understand this effect, we annotated the biological functions of the affected genes using DAVID (Huang, Sherman et al. 2007).

**Figure 2:**
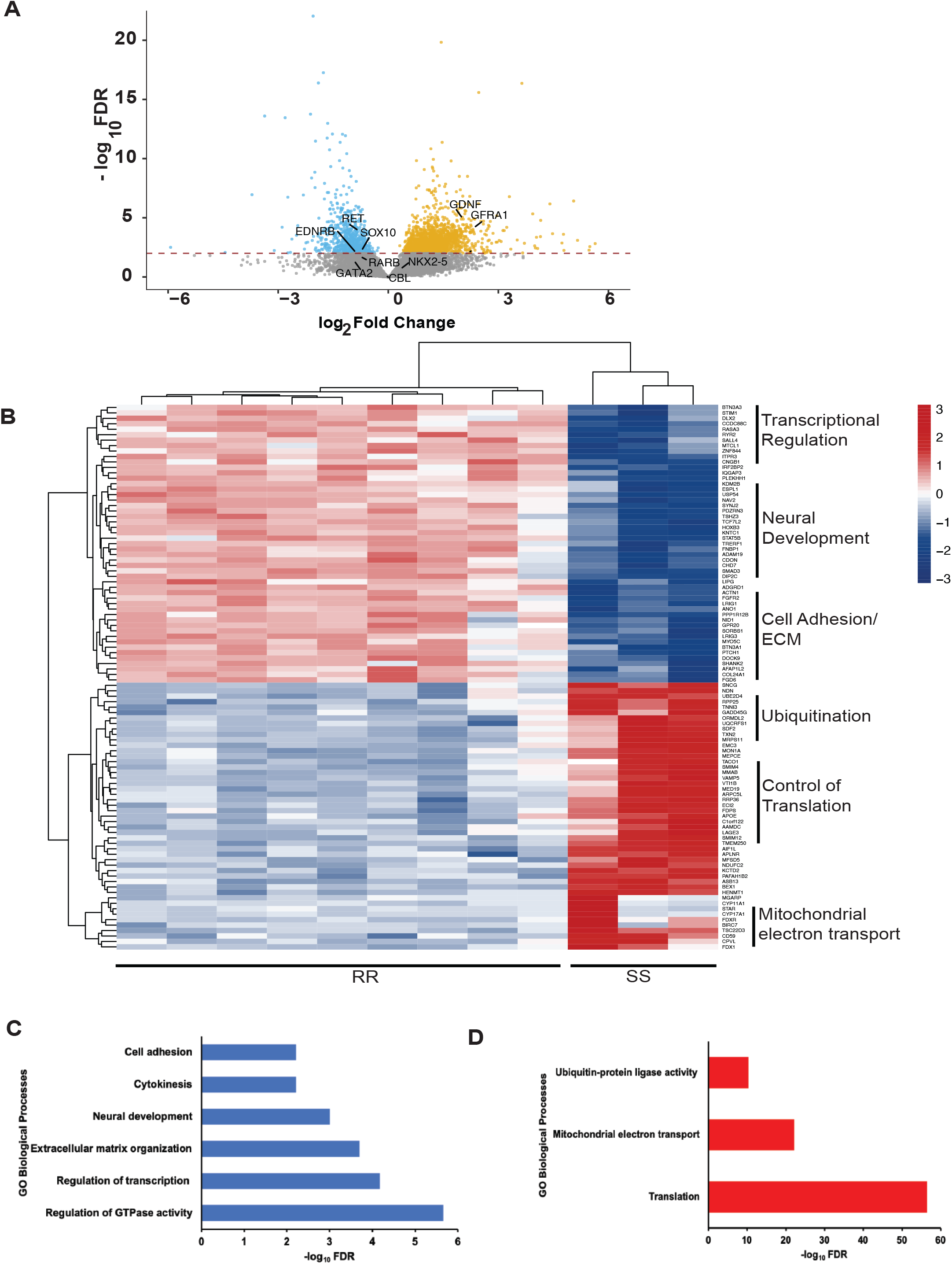
Global effects of *RET* haplotype-dependent gene expression in the CS22 human fetal gut. (A) Volcano plot of individual genes in RR versus SS embryonic CS22 guts shows 2,382 (FDR<0.01, red dotted line) differentially expressed genes with genes of the *RET-EDNRB* gene regulatory network (GRN) identified. (B) Heatmap of the top 100 genes differentially expressed between RR and SS gut tissues. (C) 1,655 genes downregulated in SS are enriched for 6 biological processes: regulation of GTPase activity, transcriptional regulation, extra cellular matrix organization, neural development, cytokinesis and cell adhesion. (D) 727 genes upregulated in SS are enriched for 3 biological processes: protein translation, mitochondrial electron transport and ubiquitin-protein ligase activity

The 1,655 downregulated genes belong to 3 significantly enriched (FDR<0.01) groups: regulatory functions, tissue morphogenesis and cell division, and motility (**Figure 2B, C**). The regulatory functions themselves are of two distinct types: regulation of transcription, involving major TFs like *PHOX2B, SALL4*, and *SOX10*, and regulation of GTPase activity with genes like *ARHGAP11A, DOCK10*, and *RASGEF1B*. These downregulated genes had largely neuronal and ECM functions. RET’s prominent effect on neuronal tissue is expected given its critical role in differentiation of ENCDCs to enteric neurons (Heanue and Pachnis 2007). This group of affected genes contains known neuronal markers like *ASCL1, NF1*, and *PLXNA2* etc. It is noteworthy that PLXNA2 is a receptor for SEMA3A, whose role in in the migration of enteric neural crest cells is well documented (Anderson, Bergner et al. 2007, Gonzales, Le Berre-Scoul et al. 2020). Moreover, multiple members of the Semaphorin class 3 genes, including *SEMA3A*, harbor pathogenic variants in HSCR as well (Luzon-Toro, Fernandez et al. 2013, Jiang, Arnold et al. 2015). Thus, three critical signaling pathways in ENS development, with receptors on the ENCDC cell surface, RET, EDNRB and class 3 Semaphorins, can transcriptionally affect each other. Decreased *RET* expression also has a significant effect on the ECM, via the reduced expression of genes like *LAMA1, ADAMTS20*, and *COL5A1* (**Figure 2C**). We have previously shown non-cell autonomous effects of RET in mouse models of aganglionosis (Chatterjee, Nandakumar et al. 2019), but these new data imply a much larger role of RET during gut morphogenesis, beyond the differentiation and proliferation of ENCDCs. We also observed additional significant effects in SS genotypes (**Figure 2C**): reduced expression of kinesin family protein genes like *KIF23, KIF20A*, and *INCENP*, hinting at cell division defects, as well as effects on cell adhesion genes like *PCDH18, LAMA3*, and *CDH13*. HSCR is characterized by loss of ENCDC cell proliferation and migration, but these data suggest that they may also involve defects in cell division and disruptions in the ECM through which ENCDCs migrate.

Genes upregulated in SS embryos also belong to three major classes (**Figure 2D**), the largest of which is protein translational control with multiple ribosomal protein genes like *RPL5, RPL30*, and *MRPS17*. We also observe an up-regulation of *NDUFB9, NDUFA2*, and *COQ9* genes, all involved in mitochondrial electron transport. Further, we see upregulation of general ubiquitination pathway genes including *UBB, UBA52*, and *PSMD13*. Perhaps, this is the leading mechanism for protein degradation in ENCDCs, since our prior studies suggest a specific connection between ubiquitination, *RET* expression and HSCR. First, *Ret* deficient mouse models have reduced expression of *Cbl*, the specific ubiquitin ligase for Ret, in the developing mouse gut (Chatterjee, Kapoor et al. 2016). Second, we have identified an enrichment of pathogenic variants in HSCR in *UBR4*, another ubiquitin ligase gene, knockdown of which leads to loss of enteric neurons in zebrafish embryos (Tilghman, Ling et al. 2019). Thus, other ubiquitin ligase complex genes may turn out to be intrinsic players in HSCR.

### RET-deficiency reduces ENCDC proliferation

The magnitude and diversity of genetic changes arising from *RET* deficiency suggests to us a parsimonious hypothesis for these observations since RET plays a direct role in the proliferation of the ENCDCs prior to enteric neuron differentiation (Vincent, Chatterjee et al. 2021). We hypothesize that quantitative decreases in *RET* ENCDC numbers, thereby reducing expression of ENCDC-expressed genes, while consequently leading to a relative increase in the proportion of non-ENCDC cells (increasing expression of genes in this cell population). This hypothesis is supported by our single cell expression study on purified *Ret* expressing ENCDC cells in the developing mouse gastrointestinal tract (Vincent, Chatterjee et al. 2021). Gene set enrichment analysis of differentially expressed genes between wildtype and *Ret* deficient cells identified several cell cycle-associated gene sets including mitotic cell cycle processes, cell division, DNA replication, regulation of cell cycle as top scoring Gene Ontology Biological Processes. Furthermore, immune-fluorescence assays on the mitotic marker phosphohistone H3 (pH3) demonstrated that there is overall reduction of pH3-marked *Ret* null cells indicating that a smaller fraction of these ENCDC cells are actively engaging the cell cycle in the absence of *Ret* leading to an overall reduction of these cells in the developing mouse gut (Vincent, Chatterjee et al. 2021).

To test if *RET* LoF leads to reduced ENCDCs in the developing human gut, we used data from the recent single cell RNA-seq study of 62,849 cells isolated from 6-11 weeks post-conception (wpc) of the developing human gut, including intestinal cells from the duo-jejunum, ileum and colon (Elmentaite, Kumasaka et al. 2021) (https://www.gutcellatlas.org). These authors identified 21 cell types including different types of epithelium and mesoderm-derived (primarily smooth muscle) tissues (**Supplementary Figure 1**). These include 4,965 cells labeled as enteric neurons or neural crest cells, which cluster together highlighting their common origin and function at this stage in development. Given the developmental stages studied (6-11 weeks wpc) it is likely that the cells labelled as neural crest cells have differentiated towards a more mature enteric neuronal identity rather than retaining their multipotency. Hence, we considered both of these clusters as ENCDCs. These cells represent ∼8% of the total cellular landscape of the GI tract at this developmental time and is similar to the 5-8% estimates in other rodent species (Hao and Young 2009, Furness 2012). We labelled all other cell types as non-ENCDC cells (57,884 cells).

We used these data to detect (>0 unique molecular identifiers) 16,013 and 17,398 protein-coding RefSeq genes expressed in the ENCDC and non-ENCDC cells, respectively. Next, we conducted differential gene expression analyses using the FindMarkers feature in *Seurat* (Stuart, Butler et al. 2019) between these clusters to detect 488 genes with significantly higher expression in ENCDC and 4,276 genes with significantly higher expression in non-ENCDC cells (FDR<1%). We then considered those genes with >2-fold significant differences in expression between RR and SS at CS22. We discovered that among downregulated genes in SS, 15% (54 of 363) were ENS-enriched while among upregulated genes in SS, only 2% (7 of 368) were ENS-enriched, a highly significant difference (P<10^−5^; Fisher’s exact test). These results can simply be explained by ENCDC cell autonomous effects from *RET* deficiency. Alternatively, more complex non-ENCDC non-cell autonomous effects could be invoked to explain down-regulated genes, however, by parsimony, the simplest hypothesis is one of ENCDC cell loss from *RET* deficiency.

These data also allow us to estimate the degree of cell loss. To do so, we identified 43 genes that have a ≥100-fold excess of ENCDC versus non-ENCDC expression and estimated the fraction of SS gene expression relative to RR gene expression from our bulk RNA-seq data (**Supplementary Table 1**). Of these, 29 were statistically significant, including the HSCR genes *RET, SOX10* and *LICAM*, which were 27%, 74% and 69% of RR expression in the SS guts, respectively. These results suggest that the population of *RET* positive ENCDCs are reduced by 73%; in contrast, *SOX10* and *LICAM* positive ENCDCs are less reduced by 26% and 31%, respectively. This suggests that transcriptionally distinct populations of ENCDCs may be differentially lost in the SS developing gut but that other related ENCDCs are less affected.

### The changing gene expression landscape during fetal gut development

To understand temporal gut morphogenesis through gene expression, and its role in HSCR, we next performed differential gene expression analysis between CS14 and CS22, the stages marking the beginning and end of gut neurogenesis (Goldstein, Hofstra et al. 2013). Because RS haplotype guts have near identical gene expression patterns as the “wildtype” RR haplotype (see before), we combined samples with these two haplotypes to compare 3 samples at CS14 with 17 non-SS samples at CS22. There were 657 and 991 genes with significantly (FDR<0.001) greater and lesser expression at CS14 versus CS22 (**Supplementary Figure 2A**). Functional annotation analysis using DAVID (Huang, Sherman et al. 2007) highlights that there is a significant enrichment of 3 classes of genes at CS14: (1) TFs, (2) genes controlling neuronal development and migration, and (3) genes in the WNT signaling pathway (**Supplementary Figure 2B**). Among these, TFs at CS14 are classic bHLH TFs like *NEUROG2* and *NEUROD4* which contribute to neuronal identity and fate commitment (Aydin, Kakumanu et al. 2019, Dennis, Han et al. 2019) and enteric neural crest TF like *PAX3* (Lang, Chen et al. 2000, Bondurand, Natarajan et al. 2006) which regulates *RET* (Chatterjee, Karasaki et al. 2021). Additionally, TFs which specify neural crest cell borders like *TFAP2* and *ZIC1* (Meulemans and Bronner-Fraser 2004) are also highly expressed in early gut development, highlighting the transcriptional memory of being derived from neural crest cells.

We next compared these data to our prior studies of gene expression in the *Ret* wildtype mouse developing gut where 89 TFs were enriched in the early stages (E10.5) (Chatterjee, Nandakumar et al. 2019). We discovered that 23 out of the 89 (27%) TFs are highly expressed both at mouse E10.5 and human CS14 gut. These evolutionary conserved TFs (*FOXC1, SALL1, LEF1, E2F5, HOXB8, MSX1, LIN28B, EMX2, LIN28A, EBF3, ALX4, HOXB9, FOXC2, POU4F1, TWIST1, PAX3, ALX3, PRRX2, SIX1, MSX2, IRX3, HOXC9 and PRRX1*) are broadly expressed in many tissues and single cell RNA-seq studies have demonstrated that a subset of these (*SALL1, LEF1, HOXB8, MSX1, TWIST1, PAX3, ALX3, PRRX2, SIX1 and IRX3*) are specifically expressed in enteric neurons of the colon in the mouse embryo and adult human (Lasrado, Boesmans et al. 2017, Drokhlyansky, Smillie et al. 2020). Thus, beyond the previously identified TFs with clear roles in HSCR, like *SOX10, PAX3, GATA2, RARB* and *NKX2-5* (Lang, Chen et al. 2000, Bondurand, Natarajan et al. 2006, Chatterjee and Chakravarti 2019), there are at least 10 additional TFs which help initiate the process of differentiation of enteric neural crest cells into enteric neurons in both mice and humans. These are attractive targets for mutation studies in HSCR.

Among other classes of early expressed genes, neuronal migration genes like *NTRK2, GJA1*, and *RELN* are enriched (**Supplementary Figure 2B**). This enrichment of ENS TF and migration genes shows that initiation of neurogenesis is one of the central processes in early gut development. The third category of early enriched genes are members of the WNT signaling pathway (e.g., *WNT9B, WNT7A*, and *DRAXIN*). The previously described TFs *LEF1* and *PITX2* are also part of the Wnt-mediated beta-catenin signaling regulatory cascade, providing additional evidence of activation of this pathway in early gut development. Conversely genes with significantly higher expression at CS22 versus CS14 fall into 4 significant (FDR<0.001) biological categories: (1) cell adhesion, (2) synaptic transmission, (3) smooth muscle contraction and (4) MAP kinase signaling (**Supplementary Figure 2C)**. This follows the pattern observed in mouse gut development in that the later stages are dominated by the emergence of specialized structures and new functions (Chatterjee, Nandakumar et al. 2019), such as cell adhesion (*CNTNAP1, CNTNAP2* and *COL16A1*), synaptic transmission (*KCNA1, NRXN1* and *HTR2B*), and, smooth muscle contraction (*SMTN, MYH11* and *MYLK*); we also see an activation of multiple genes in MAP kinase signaling (*TENM1, FLT3* and *KITLG*). Thus, the later stages of gut development are marked with increased activity of genes helping in the formation of complex tissues and cellular communication via many signaling pathways.

### The transcriptional repertoire controlling RET and EDNRB gene expression

To ascertain how many of the gut developmental TFs we discovered control *RET* and *EDNRB*, we performed siRNA-mediated knockdown of each in the human neuroblastoma cell line SK-N-SH (Chatterjee, Kapoor et al. 2016, Chatterjee and Chakravarti 2019). We utilized our larger expression dataset of the mouse gut development in wildtype and *Ret* deficient guts (Chatterjee, Nandakumar et al. 2019) to select two classes of TFs affected by *Ret* LoF: (1) 25 TFs which show ≥2-fold expression at E10.5 versus E14.5 (*early Ret TFs*), and (2) 14 TFs common throughout gut development (*common Ret TFs*) (**Table 2**).

**Table 2:** List of 25 transcription factors (TFs) affected by *Ret* deficiency in the early mouse embryonic gut and 14 TFs all through development (common TFs). TFs in bold are not expressed in SK-N-SH cells and were excluded from our knockdown screen.

| Early *Ret* controlled TFs | Common *Ret* controlled TFs |
| --- | --- |
| **MYOG** | *ASCL1* |
| *LHX9* | *TBX2* |
| **PRRX1** | *BARX1* |
| *BCL11A* | *SOX10* |
| **HOXB9** | *BAZ2B* |
| *TWIST1* | *SNAI1* |
| *FOXC1* | *ZFHX4* |
| *FOXD1* | *PLAG1* |
| *ISL1* | *BNC2* |
| *HOXC9* | *TBX3* |
| *HOXC8* | **FOXP2** |
| *HOXC6* | *FOXP1* |
| *BACH1* | *PEG3* |
| *SIX2* | *NFAT5* |
| **ZFP521** |  |
| *EMX2* |  |
| *PRRX2* |  |
| *ALX4* |  |
| *SALL4* |  |
| *LEF1* |  |
| **HOXA9** |  |
| *GATA2* |  |
| *EBF3* |  |
| *SALL1* |  |

We first searched the expression profile of all 39 TFs in a published SK-N-SH RNA-seq data (Harenza, Diamond et al. 2017). Of these, we did not detect expression of *MYOG, PRRX1, HOXA9, HOXB9, ZFP521* and *FOXP2* in this human cell line (**Table 2**), and so they were not studied further. As a positive control we first knocked down *RET* and *EDNRB* to demonstrate that their respective gene expression decreased to 26% (P = 4.1 × 10^−8^) and 15% (P = 6.4×10^−6^) (**Figure 3A, 3B**), as compared to control siRNAs, recapitulating our prior data (Chatterjee and Chakravarti 2019). We also demonstrated transcriptional feedback between *RET* and *EDNRB* by observing a 22% decrease (P = 4×10^−4^) in *EDNRB* expression from *RET* knockdown, and a 34% decrease (P = 3.2×10^−5^) in *RET* expression from *EDNRB* knockdown (**Figure 3A, 3B**).

**Figure 3:**
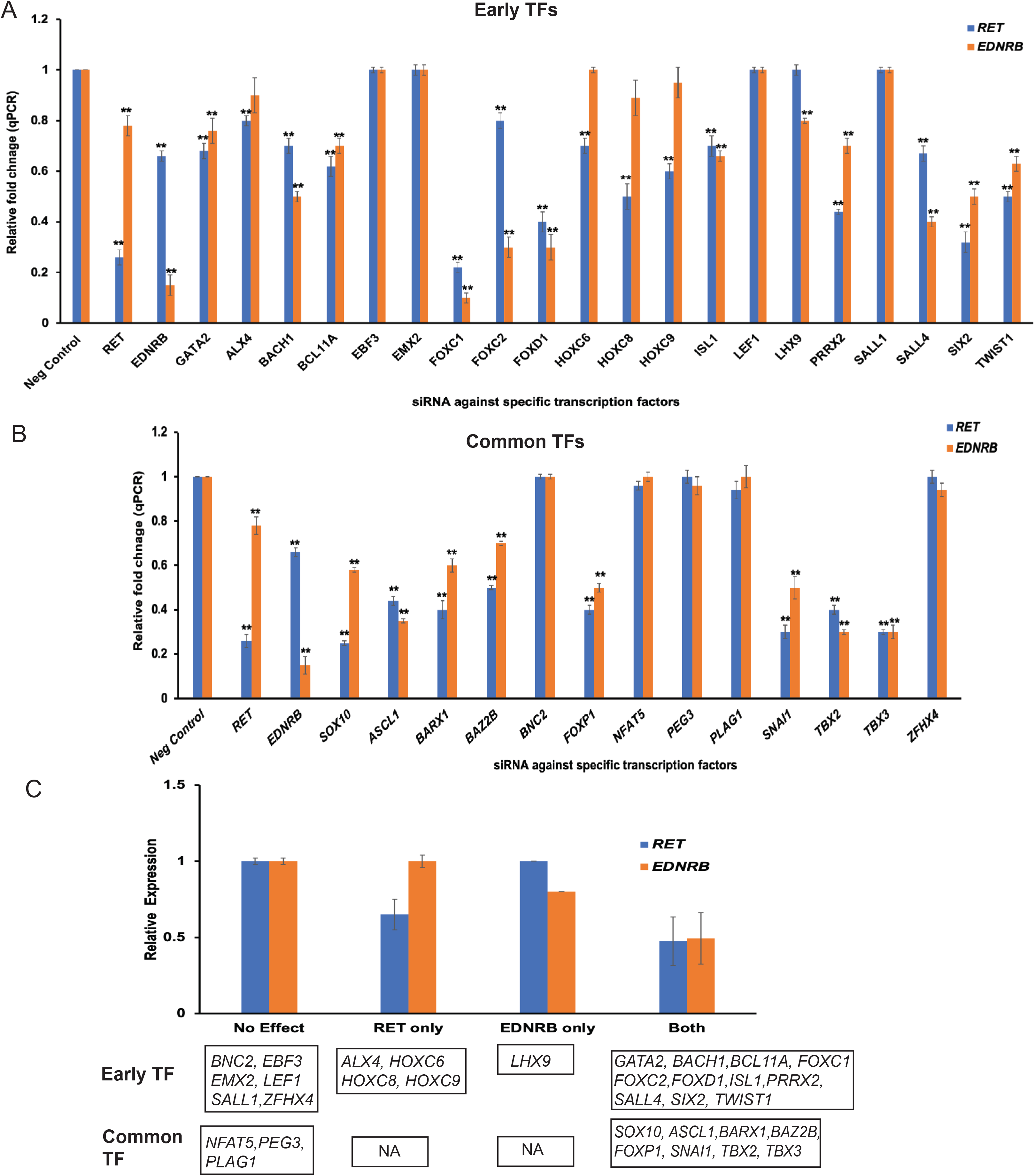
Transcription factors controlling ENS development. (A) siRNA knockdown in the SK-N-SH cell line for 20 transcription factors (TFs) that show enriched expression in early (CS14) human fetal gut development and whose expression is affected by *RET* deficiency. Of these, 11 TFs significantly (P <0.001) alter *RET* and *EDNRB* expression, 4 alter *RET* expression only, and 1 affected *EDNRB* expression only. A similar analysis of 13 TFs whose expression pattern is constant throughout development and also affected by *RET* deficiency identified 8 TFs that significantly (P <0.001) alter both *RET* and *EDNRB* expression. (C) This analysis identified 24 TFs with bidirectional transcriptional feedback with *RET* and/or *EDNRB*, of which 19 are potential HSCR genes.

Next, we performed siRNA-mediated knockdown of the 20 early mouse TFs expressed in our cell line, 12 of which are also expressed at a higher level in CS14 versus CS22, to demonstrate that 11 significantly (P <0.001) altered *RET* and *EDNRB* expression, 4 affected *RET* only and 1 affected *EDNRB* only (**Figure 3A**). Of 13 TFs affected throughout gut development, upon perturbation, we identified 8 TFs, namely, *ASCL1, BAZ2B, BARX1, FOXP1, SNAI1, SOX10, TBX5*, and *TBX3*, that significantly (P <0.001) altered both *RET* and *EDNRB* expression (**Figure 3B**). Thus, these 24 TFs are part of an exte*nded RET-EDNRB GRN*, of which 19 TFs affect the gene expression of both *RET* and *EDNRB* (**Figure 3C**). These genes are prime candidates for mutation screening in HSCR patients.

## Discussion

One of the major physiological functions of the gut is motility arising from an extensive neural network in the gut. Formation of this enteric nervous system occurs through differentiation of enteric neural crest derived cells to enteric neurons, a key developmental milestone of fetal gut morphogenesis (Burns and Douarin 1998). One of the major genes in this fate transition is *RET*, which harbors coding and regulatory pathogenic alleles (PAs) in ∼50% of Hirschsprung disease (HSCR) cases (Tilghman, Ling et al. 2019). Curiously, the majority of this risk arises from a single susceptibility haplotype (S) containing three hypomorphic enhancers of *RET*. Our more recent studies, have identified an additional 7 functional *RET* enhancers each with a HSCR-associated variant and extended the molecular dissection of this risk haplotype (Chatterjee, Karasaki et al. 2021). These data suggest that the high risk of this regulatory haplotype, resulting in 3-fold decreased *RET* gene expression (**Figure 1B**), likely arises from the multiple enhancer defects it harbors.

The impact of the SS homozygote on its developing gut transcriptome is substantial, affecting 8% of all genes across diverse pathways, specifically genes expressed in the ENCDC cell population. Our results strongly suggest that this large effect arises from substantial loss of proliferation of ENCDC cells with a concomitant increase of non-ENCDC cells. This loss is as large as 73% in RET-positive cells although other subpopulations of ENCDCs have attenuated effects. Thus, the SS gut has a different cell type distribution than the RR gut, likely altering its neuronal biology. In HSCR this cell loss is even more severe, resulting in aganglionosis, altering its biology to a pathology, and involving all cell types along the ENCD – non-ENCDC axis. *RET* deficiency induces major cell autonomous and non-cell autonomous, primarily in smooth muscle and epithelium development, effects we have previously observed in *Ret*-deficient mouse models (Chatterjee, Nandakumar et al. 2019). These non-neuronal genetic changes may be a potential explanation of why ∼32% of HSCR patients have developmental gut anomalies (e.g., malrotation of the gut) beyond aganglionosis (Tilghman, Ling et al. 2019). Thus, HSCR is best viewed as a multifactorial disorder of the gut involving the pathophysiology of an altered gut cell distribution, implications that are testable in human gut surgical resection samples.

The cell loss in the *RET* deficient developing gut raises the question of whether the transcriptomic changes are from changes in gut cell composition only or also from regulatory alterations within the *RET-EDNRB* GRN. We believe the latter because *RET* deficiency alters the levels of biochemically proven TFs, ligands and E3 ligases. Our experimental analyses clearly show that 24 ENCDC-enriched TFs are direct regulators of *RET* and/or *EDNRB* gene expression, and are, consequently, a part of the *RET-EDNRB* GRN. Of these, at least 8 are significantly different in the RR versus SS gut (**Figure 2A**).

The final enigma of the RR versus SS comparisons is that despite the extensive genetic and cell type changes we observed, HSCR aganglionosis is much rarer than the SS genotype: why? The latter probably occurs through additional genetic and/or epigenetic changes. If so, this study has provided many functional candidate genes that could be specifically screened for HSCR pathogenic variants. An interesting direction would be to compare, by genome sequencing, gene expression and methylation assays, gut tissue from HSCR affected and unaffected SS individuals.

## Materials and Methods

### Human fetal gut tissues

Frozen human fetal gut tissues were obtained from the Human Developmental Biology Resource (www.hdbr.org), voluntarily donated by women undergoing pregnancy termination following specific written consent. All samples are anonymous. The tissues are karyotyped and samples with large chromosomal aberrations removed from the collection. These studies were conducted with approval by the Institutional Review Boards of NYU Grossman School of Medicine (s17-01813). For these studies, we received 23 tissue samples.

### Genotyping

We genotyped the three *RET* enhancer single nucleotide polymorphisms (SNPs) rs2506030, rs7069590 and rs2435357 using specific TaqMan Human Pre-Designed genotyping assays following the manufacturer’s protocol (ThermoFisher Scientific). The assays IDs are C_26742714_10, C_2046272_10 and C_16017524_10 for rs2506030, rs7069590 and rs2435357, respectively. The end-point fluorescence measurements were performed on a 7900HT Fast Real-Time PCR System (Applied Biosystems) and analyzed using Sequence Detection System Software v.2.1 (Applied Biosystems).

### RNA extraction and sequencing

Total RNA was extracted from each sample using TRIzol (Life Technologies, USA) and cleaned on RNeasy columns (Qiagen, USA). Sample integrity (>9 RIN) was assessed using an Agilent 2100 Bioanalyzer (AgilentTechnologies) and cDNA prepared using oligo dT beads to select mRNA from total RNA followed by heat fragmentation and cDNA synthesis, as part of the Illumina RNA Sample Preparation protocol. The resultant cDNA was then used for library preparation (end repair, base ‘A’ addition, adapter ligation, and enrichment) using standard Illumina protocols. Libraries were run on a HiSeq 2500 instrument to a depth of 50 million reads per samples (paired-end 100 base pair reads), using the manufacturer’s protocols. The BCL (base calls) binary files were converted into FASTQ using the bcl2fastq Illumina package. RNA-seq paired-end read fastq files were quality checked using FASTQC and then processed using Trimmomatic (Bolger, Lohse et al. 2014) for removing adapters and other Illumina specific sequences from the reads, and for performing a sliding-window based trimming of low quality bases from each read (ILLUMINACLIP:TruSeq3-PE-2.fa:2:30:10:1:TRUE LEADING:3TRAILING:3 SLIDINGWINDOW:4:15 MINLEN:36). RNA Read alignment was performed using the STAR 2.6.0a software (Dobin, Davis et al. 2013) against the human reference genome hg19. During alignment, non-canonical junctions was removed (option: --outFilterIntronMotifs RemoveNoncanonical), XS strand attributes were generated for splice alignments (option: --outSAMstrandField intronMotif), and the numbers of reads per gene were counted (option: --quantMode GeneCounts). Next, differential expression analysis was conducted using DESeq2 (version: 1.24.0) (Anders and Huber 2010) with default parameters. Significant differentially expressed genes (DEG) was defined as any gene with Benjamini-Hochberg adjusted P ≤ 0.01, absolute fold change ≥ 2, and read count ≥ 5 in at least one group. All raw reads files and normalized read counts have been uploaded to the Gene Expression Omnibus (GEO) under accession number GSE160359.

### siRNA assays

Silencer select siRNA library, a combination of 3 individual siRNAs targeting each gene, obtained from NYU Langone’s High Throughput Biology Laboratory, along with a non-targeting negative control siRNA (**Supplementary Table 2**) were transfected at 25 µM concentration in SK-N-SH cells at a density of 10^4^ -10^5^ cells using FuGene HD Transfection reagent (Promega Corporation, USA), per the manufacturer’s instructions. At a 25 µM concentration, the expression level of each gene was below 50% of its expression when compared to the negative control siRNA.

Total RNA was extracted from cells 48-hour post-transfection and Taqman gene-specific assays conducted as described. Three independent transfections were used for each siRNA and each Taqman assay was performed in triplicate (n = 9); P values were calculated from pairwise 2-tailed t-tests and the data presented as means with their standard errors (SE).

### Gene expression Taqman assays

Total RNA was extracted from SK-N-SH cells using TRIzol (Life Technologies, USA) and cleaned on RNeasy columns (QIAGEN, USA). 300µg of total RNA was converted to cDNA using SuperScriptIII reverse transcriptase (Life Technologies, USA) using Oligo-dT primers. The diluted (1/5) total cDNA was subjected to Taqman gene expression analyses (ThermoFisher Scientific) using transcript-specific probes and primers (**Supplementary Table 3**). Human β-actin was used as an internal loading control for normalization. Three independent wells for SK-N-SH cells were used for RNA extraction and each assay was performed in triplicate (n = 9). Relative fold change was calculated based on the 2ΔΔCt (threshold cycle) method. For siRNA experiments, 2ΔΔCt for negative control non-targeting control siRNA was set to unity. P values were calculated from pairwise 2-tailed t-tests and the data presented as means with their standard errors (SE).

### Single cell RNA-seq data

The processed AnnData file for 62,849 cells isolated from 6-11 weeks post-conception developing human gut was downloaded from https://www.gutcellatlas.org/ (Elmentaite, Kumasaka et al. 2021). This was converted to a Seurat object using the “Convert” function in *Seurat* (Stuart, Butler et al. 2019). We retained all the cells from the original data did not perform any further quality control steps to remove cells based on gene and UMI count or mitochondrial reads. We used the authors described 10 Principal components to generates the Uniform Manifold Approximation and Projection (UMAP) embedding on these cells using the python package “umap-learn” in *Seurat*. The cell identity of the UMAP clusters was derived from the metadata file provided by the authors. We considered the cells identified as neural crest cells and enteric neurons a s single cluster called enteric neural crest derived cells (ENCDC). Differential gene expression between ENCDC cluster and all other cells was performed using “FindMarker” function in *Seurat* between protein coding genes which were detected in the cells (UMI>0). Genes significantly (FDR<0.01) expressed highly in ENCDC cluster were labelled as ENCDC enriched genes and conversely genes significantly highly expressed in the other clusters combined were labeled as non ENCDC enriched genes. Owing to the weaker detection of transcripts in standard single cell RNA-seq experiments we took a lower significance threshold to include more genes in our analysis.

## Author Contributions

A.C and S.C. conceived and designed the study. S.C., H.B-R and A.C. coordinated tissue collection from the Human Developmental Biology Resource (HDBR); S.C., H.B-R and L.E.F performed genotyping, RNA extractions and analysis; S.C., O.Y and N.H. conducted expression analyses; S.C. and A.C. wrote the manuscript.

## Supplementary Table legends

**Supplementary Table 1:** 43 RefSeq protein coding genes which have >100-fold expression in enteric neural crest derived cells as compared to non-enteric neural crest derived cells together with their expression in RR and SS embryonic guts.

**Supplementary Table 2:** Sequences of siRNAs used to knockdown transcription factors in the SK-N-SH cell line.

**Supplementary Table 3:** Assay IDs for Taqman qPCR probes and primers.

## Supplementary Figure legends

**Supplementary Figure 1: Cellular consequences of *RET* loss of function (LoF) in the developing human fetal gut**.

The diversity and distribution of 62,849 cells (UMAP, Uniform Manifold Approximation and Projection) in the developing human fetal gut at 6-11 weeks post-conception identifies 21 major cell types (Elmentaite, R et.al. *Nature* 2021). The enteric neural crest-derived cells (ENCDC) comprise 8% of the total and form 2 closely associated cell clusters.

**Supplementary Figure 2: The temporal gene expression program of the developing human fetal gut at CS14 and CS22**.

(A) Heatmap showing the top 100 differentially expressed genes at CS14 and CS22 stages of gut development. (B) At CS14, 657 genes show greater expression and are enriched for transcriptional regulation, neuronal migration and *Wnt* signaling functions, whereas (C) at CS22, 991 genes show greater expression and are enriched for organogenesis processes, such as, smooth muscle contraction, synaptic transmission along with cell adhesion and regulation of MAP kinase pathways.

