## Supplementary Figures for "*RET* enhancer haplotype-dependent remodeling of the human fetal gut development program"

### SUPPLEMENTARY FIGURE 1

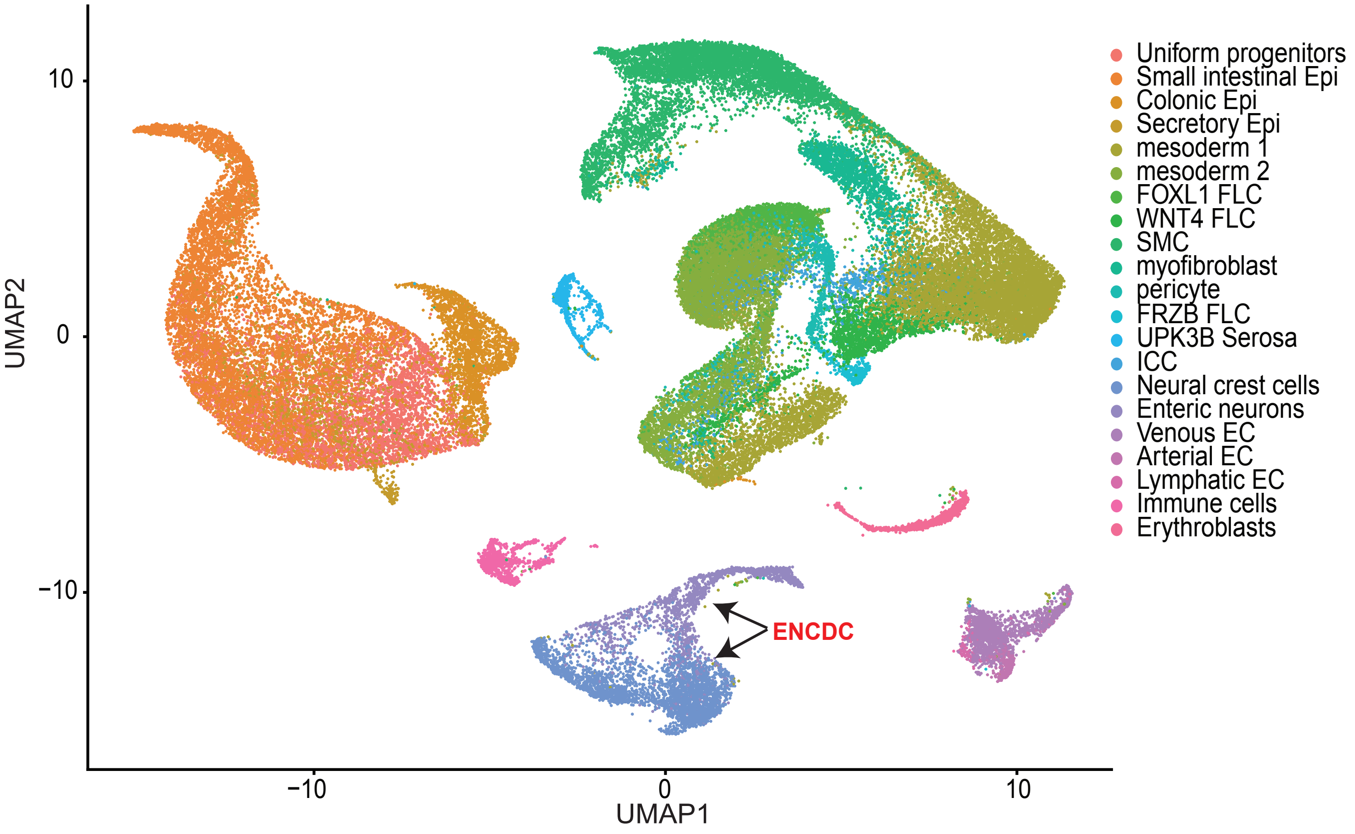

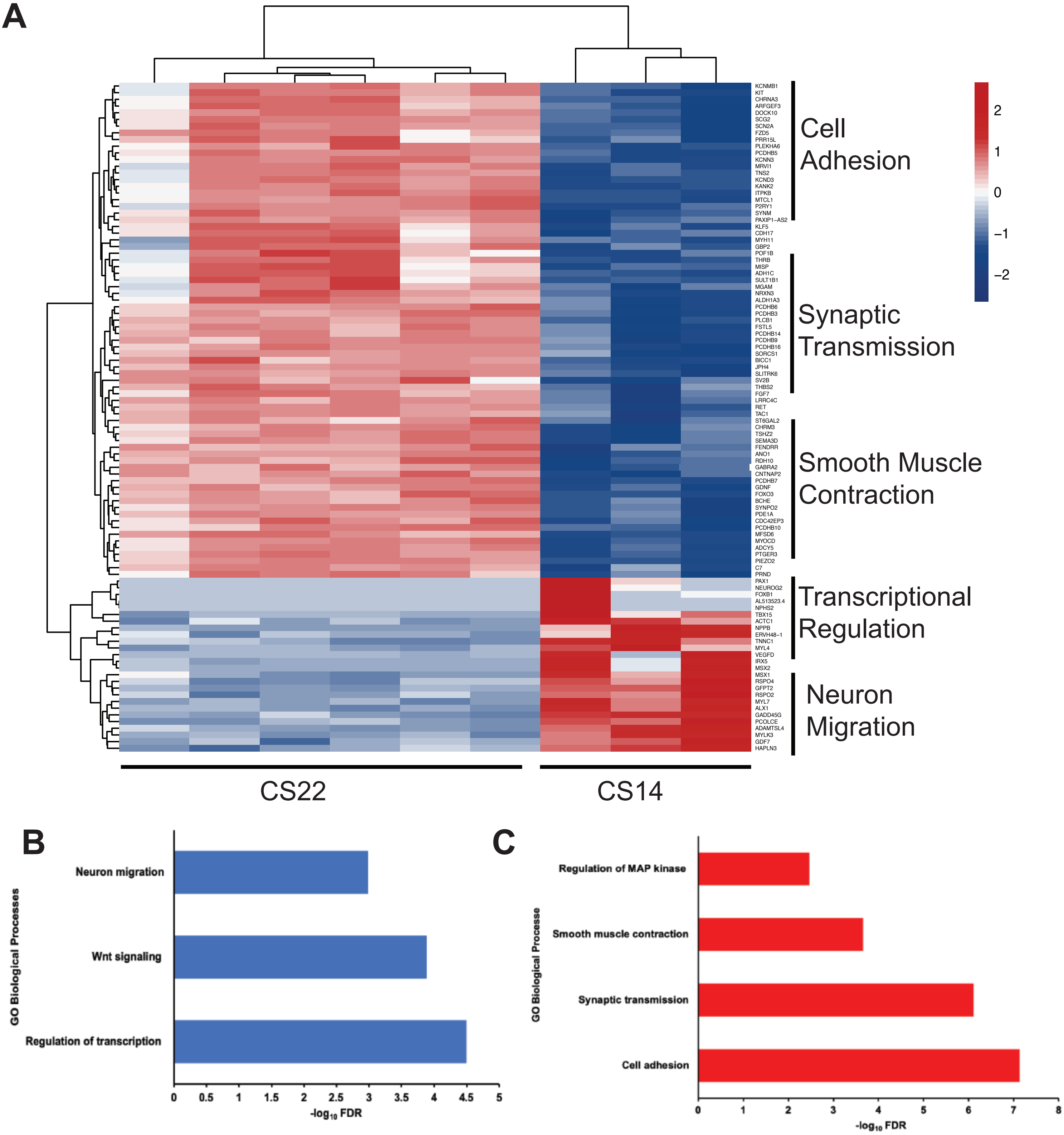


Genes upregulated at CS14
(Early GI tract genes)

Genes upregulated at CS22
(Late GI tract genes)
